# Joint representations of color and form in mouse visual cortex described by random pooling from rods and cones

**DOI:** 10.1101/2021.07.26.453648

**Authors:** I. Rhim, I. Nauhaus

**Author notes:** **Correspondence:** Ian Nauhaus, Department of Psychology, Center for Perceptual Systems, University of Texas at Austin, 108 E. Dean Keeton, Austin TX, 78712.

## Abstract

Spatial transitions in color can aid any visual perception task, and its neural representation – the “integration of color and form” – is thought to begin at primary visual cortex (V1). An integration of color and form is untested in mouse V1, yet studies show that the ventral retina provides the necessary substrate from green-sensitive rods and UV-sensitive cones. Here, we used two-photon imaging in V1 to measure spatial frequency (SF) tuning along four axes of rod and cone contrast space, including luminance and color. We first reveal that V1’s sensitivity to color is similar to luminance, yet average SF tuning is significantly shifted lowpass for color. Next, guided by linear models, we used SF tuning along all four color axes to estimate the proportion of neurons that fall into classic models of color opponency – “single-”, “double-”, and “non-opponent”. Few neurons (~6%) fit criteria for double-opponency, which are uniquely tuned for chromatic borders. Most of the population can be described as a unimodal distribution ranging from strongly single-opponent to non-opponent. Consistent with recent studies of the rodent and primate retina, our V1 data is well-described by a simple model in which ON and OFF channels to V1 sample the photoreceptor mosaic randomly.

## Introduction

A dichromatic animal has two cones that filter the visual scene through unique spectral sensitivity functions, creating two retinal images that can be compared for the detection of color and luminance contrast at each scene location. Mice are dichromats, but their two cone opsins have minimal intermixing at each location of the retina, a requirement for encoding spatial color contrast. Instead, M-opsin (peak at 508 nm) is concentrated in the dorsal retina, whereas S-opsin (peak at 360 nm) is in the ventral retina. However, rods may provide the missing ingredient for color vision in mice. Rods (peak at 500 nm) have a similar spectral sensitivity as M-opsin, and intermix with S-opsin during mesopic light adaptation in the ventral retina. Indeed, studies of the retina have shown that full-field transitions between green and UV light are signaled by circuits that subtract rod and cone inputs – i.e. “single-opponency” ^1–3^, which may be linked to behavioral discrimination of related stimuli ^4^. Furthermore, spatial gradations of UV-green contrast are enriched in the upper visual field at twilight ^5^, which may be particularly valuable for aerial predator detection ^6–8^. Motivated by these recent studies, we designed experiments to probe the interplay of color and spatial contrast in the upper visual field representation of mouse V1.

Selectivity for the spatial form in an image is considerably sharpened at the level of primary visual cortex (V1) ^9–11^. Perhaps most notably, the V1 population is diversely tuned for the spatial frequency (SF) and orientation of sinewave gratings. An important question is whether the exquisite tuning for spatial form in V1 is limited to luminance contrast, or if V1 also encodes form via color contrast ^12^. Several studies in primate V1 have addressed this question by measuring spatial tuning properties with different types of color contrast ^13–16^, yielding variable results. A handle on this seemingly complex problem starts with our pre-existing knowledge of V1’s responses to grayscale sinewave gratings, and generalizes to a multi-dimensional color space where each grating modulates a specified combination of opsins in the retina ^12,13,15^. However, this framework had not been used to study mouse V1, so little was known about its spatial representation of color contrast, which could lead to new insights into the underlying mechanism with genetic tools.

Here, we investigated the joint representation of color and form in mouse V1, mediated by rod and cone S-opsin inputs. With two-photon imaging, we measured SF tuning along four axes of S-opsin vs. rod (“R-opsin”) contrast space: S, R, R-S, and R+S. We find that V1 is more sensitive to color (S-R) than luminance (S+R) contrast at low SFs, yet more sensitive to luminance than color at higher SFs. Based on classic models of color tuning in V1, this result implies that the average V1 RF in our data is “single-opponent”, meaning input from one opsin responds to increments in light (ON) and input from the other responds to decrements (OFF) ^12^. However, a detailed analysis - strongly guided by linear RF models - revealed that the population can be described as a continuum from “single-opponent” to “non-opponent”. The ON and OFF channels of a non-opponent cell (aka “luminance cell”) receive similar proportions of the two opsins. Finally, we show that this continuum is consistent with a model whereby the ON and OFF channels pool random quantities of the two opsins. Random pooling models have been shown to account for cone opponency in the primate retina ^17^, and may also describe much of the rod-cone opponency in mouse retina ^1^. We conclude that the integration of color and form in mouse V1 can be described as a parsimonious addendum to classic Gabor models ^9^.

## Results

### Models of rod-cone opponency as reference points in the data

For each detected neuron in mouse V1, we measured four SF tuning curves, one for each color contrast in our stimulus ensemble: R, S, R+S, and R-S. The R- and S-contrast isolate rods and cone S-opsin, respectively. The R+S contrast adds rods and cone S-opsin, whereas R-S subtracts them. This created a rich data set exhibiting several dependencies between spatial tuning and color. Prior to the presentation of the data, we first describe RF models that will be used to aid in its interpretation. We simulated four models of spatio-chromatic RFs in V1. Each RF model provides a point of reference within the measured parameter distributions shown in subsequent figures. They will also help to motivate the random wiring model at the very end of the Results.

The first three RF models – “single-opponent-A”, “single-opponent-B”, “non-opponent” – can originate from the same achromatic model of OFF and ON subfields (Fig. 1c). However, they differ in their linear decomposition into R- and S-opsin RFs. More specifically, they receive different proportions of R- and S-opsin input to their ON and OFF subfields, which leads to unique spatial tuning for R, S, and R-S contrast (but not S+R). Here, we highlight some of these distinctions between models that will aid in the classification of each recorded neuron. For one, single-opponent-A and non-opponent models have lowpass and bandpass SF tuning for S-R (color) contrast, respectively (Fig. 1f). Similarly, single-opponent-A and non-opponent models are lowpass and bandpass for R- and S-opsin isolating contrast, respectively. In general, the tuning of single-opponent-B lies somewhere between the other two (Fig. 1e). Finally, the preference for S-R over S+R contrast will be an important classification parameter. For example, single-opponent-A and single-opponent-B both have lowpass SF tuning for S-R contrast, but single-opponent-A prefers S-R over S+R.

**Figure 1:**
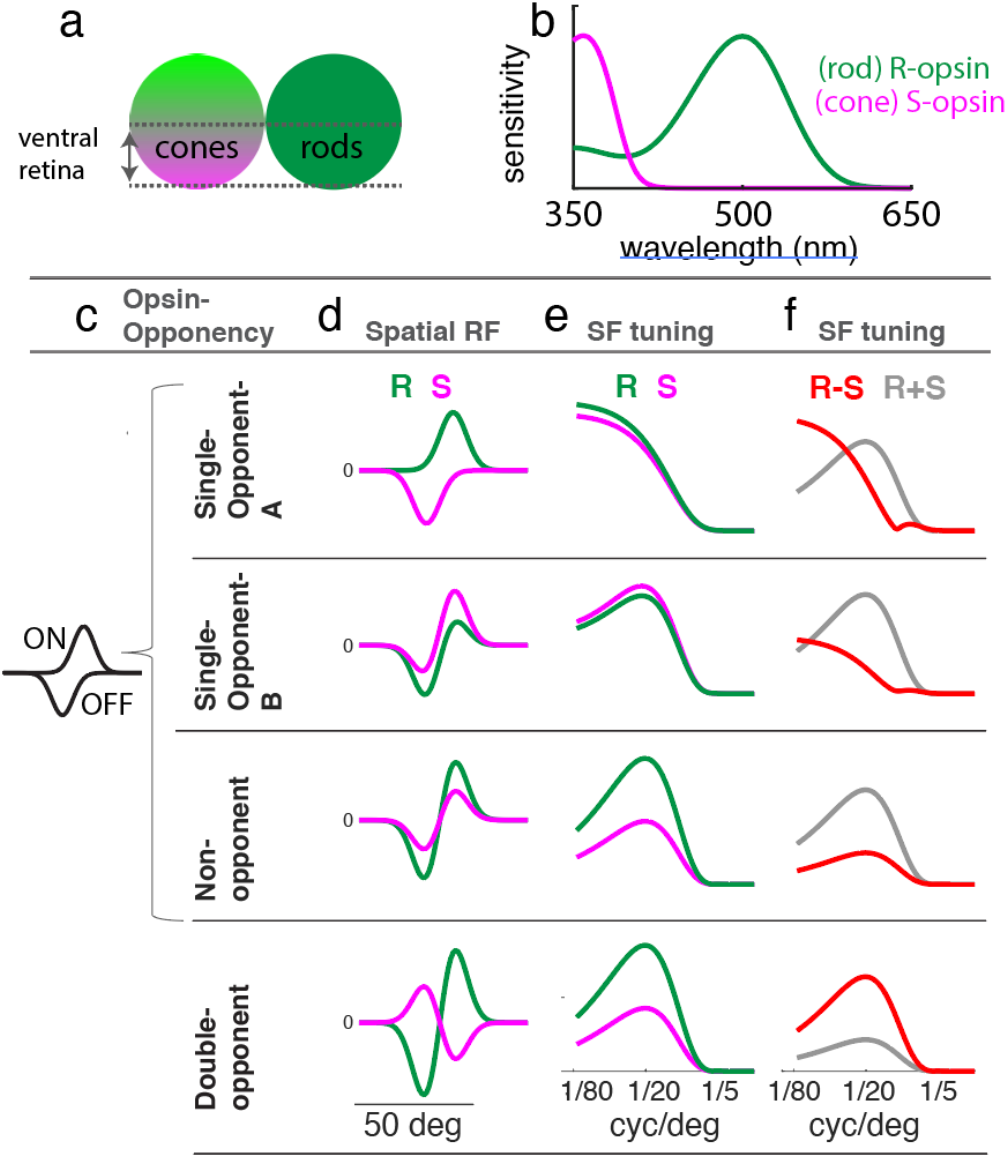
Models of rod vs. S-cone opponency. (a) Illustration of the spatial distribution of opsins in the mouse retina. In the cone mosaic on the left, green and violet indicate concentration of M- and S-opsin, respectively. The green disk on the right represents the uniform rhodopsin (“R-opsin”) distribution. (b) Approximate spectral sensitivity functions of the two major opsins in the ventral retina (Govardovskii et. al. 2000), which was the focus of this study. (c) The title in each row defines the model of color opponency. The first three are built from the same ON and OFF subfields. (d) Each row has two linear RFs, one for R-opsin (green) and one for S-opsin (violet). (e) Taking the Fourier transform of each opsin’s RF in ‘b’ approximates the sensitivity of the neuron to sinewave gratings with contrast that isolates either the R- or S-opsin axis of color space. (f) To obtain the SF tuning to R+S (black) and R-S (red) contrast, we added and subtracted the RFs in ‘d’, respectively, and then took the Fourier transform.

The fourth and final simulated model is a so-called “double-opponent” RF, which is unique from the other three in that it requires an ON and OFF subfield at each spatial location (Fig. 1d, bottom). Like the non-opponent model, SF tuning is bandpass for each axis of contrast, but with the critical distinction of being more sensitive to S-R than S+R contrast (Fig. 1f, bottom).

### Paradigm for measuring rod-cone contributions to color and form tuning in V1

To build on recent studies showing rod-cone opponency in the ventral retina ^1,2^, we biased two-photon imaging fields-of-view to the upper visual field representation of V1 (Fig. 2a,b), while the retina was in a mesopic state of light adaptation ^18^. The lack of M-opsin expression by cones in the ventral retina allowed for rod isolation, and more generally, calibrated stimuli within the R- and S-opsin contrast plane. Neuronal responses were measured with two-photon imaging of GCaMP6f in WT mice (AAV1.syn.GCaMP6f) ^19^.

**Figure 2:**
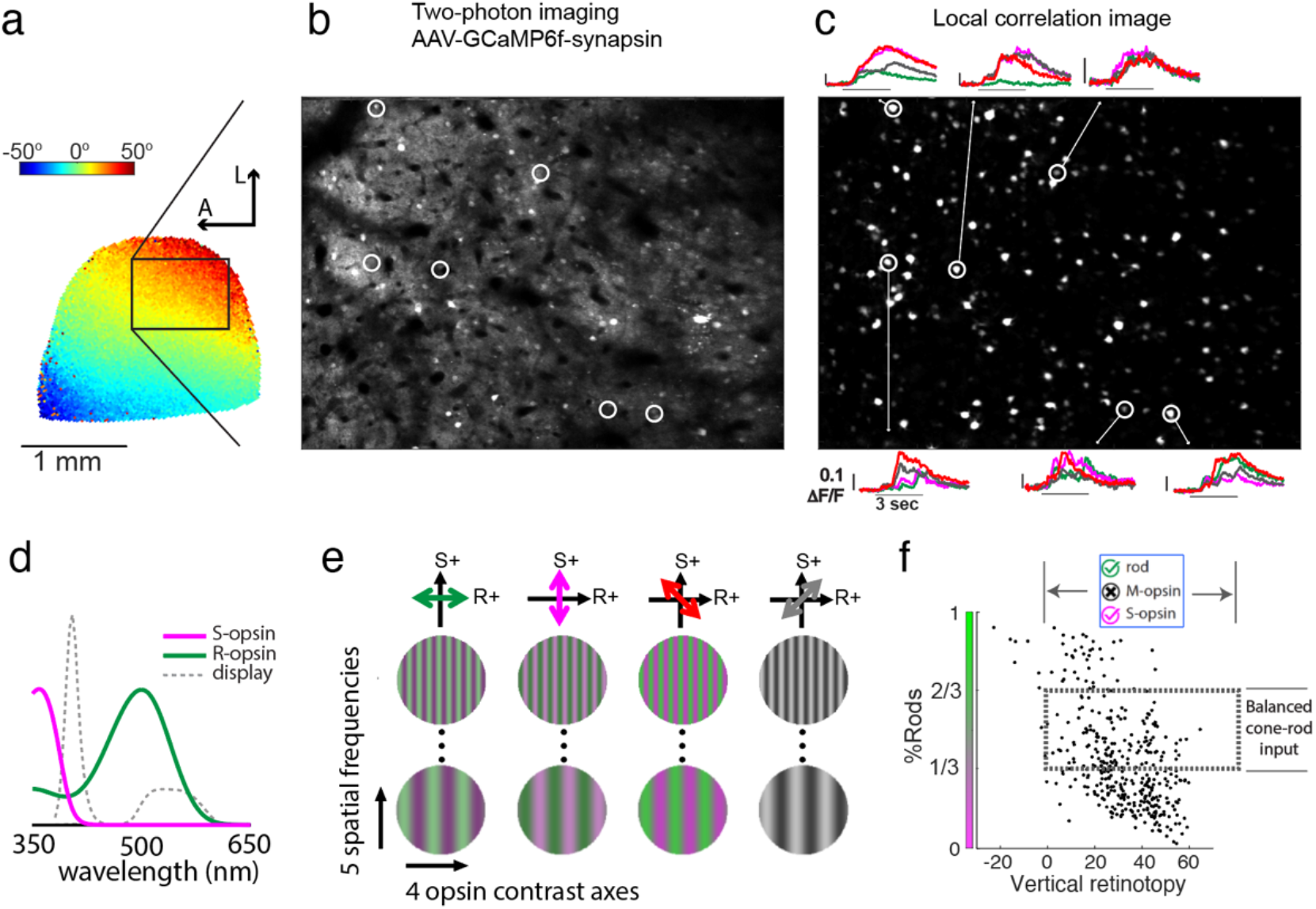
Two-photon imaging of responses to drifting gratings in the S- and R-opsin contrast plane. (a) A map of vertical retinotopy in V1 of the right hemisphere, using intrinsic signal imaging. Blue-to-red is lower-to-upper fields. Subsequent two-photon imaging was targeted to the upper visual field representation. (b) Raw image from a two-photon imaging session, 900 μm wide. (c) The local correlation image, from which 126 cell bodies were manually selected. The total across 16 ROIs using this procedure was 1360. Bright pixels are highly correlated with the immediate neighborhood. Each small panel surrounding the perimeter shows the circled neuron’s responses to different opsin contrasts, at the neuron’s optimal drift direction and SF, averaged over 5 repeats. The color indicates the axis of opsin contrast, shown in ‘e’. (d) Upper visual field opsin sensitivities (Govardovskii et. al. 2000), overlaid with the spectral power distribution of the visual stimulus. (e) Illustration of the stimulus set. There were five SFs, spaced 1 octave apart, ranging from 1/80 c/° to 1/5 c/°. In a small subset of experiments, a sixth SF was included at 1/2.5 c/°. Above each column of gratings is a diagram of the corresponding color axis. (f) Each data point is a neuron that was first manually selected from one of 16 local correlation images like in ‘c’, and then shown to produce reliable parameter estimates of SF tuning and retinotopy (see Methods), yielding n = 385 in this plot. The y-axis is the relative response to the S- and R-opsin isolating contrast. A value of 1 and 0 indicates that it only responded to R- and S-contrast, respectively. The x-axis is the RF’s position in vertical retinotopy, with positive values being above the midline. The overlaid dashed rectangle outlines the population used for subsequent analyses, which encloses neurons in the upper visual field, and neurons with a balanced R:S response ratio (1:2 < R:S < 2:1), totaling n = 140. This data selection allowed for the study of rod-cone color opponency with gratings that contain contrast in the R- and S-opsin color plane.

We measured SF tuning along four different axes in the S- and R-opsin contrast plane: R, S, S-R and S+R (Fig. 2e). Total opsin contrast was the same for each axis (60%). Figure 2c shows the responses of six example neurons, where each trace was elicited by a different opsin contrast at the optimal SF. The other stimulus used in this study was a monochrome drifting bar (not shown) to verify the upper visual field representation of each neuron ^20,21^.

A requirement for neural processing of color is simultaneous input from two types of photoreceptors. As shown in Figure 2f there is considerable variation in the balance of rod and cone input, as assessed by “%R”. %R is the response to R-contrast, divided by the summed response to S- and R-contrast. Specifically, the response to each type of contrast was defined as the peak of the respective SF tuning curve. SF tuning was the average over 5 repeats and 4 orientations. The broad distribution of %R may be due to multiple factors, such as variability in states of retinal adaptation across experiments, strength of rod vs. cone input, and trial-to-trial response. For instance, some experiments had lower %R, implying that the rods were more saturated. To proceed with an assessment of color vs. form, we selected a subpopulation of cells that were balanced in their response to the S- and R-isolating contrast: 1/3 < %R < 2/3. This translates to a response ratio between contrasts (R:S) that is somewhere between 1:2 and 2:1. In the Discussion, we summarize how key results are expected to change when the population is more imbalanced, and also re-analyze the data to validate these expectations. Finally, although each two-photon imaging region was biased to the upper field representation, the RF position of each neuron was verified. A small subset of neurons with RFs below the midline were excluded (Fig. 2f).

### V1 neurons prefer color contrast at lower SFs, and luminance contrast at higher SFs

The sensitivity of V1 to color is an important and contentious question in vision science ^22^, but has not been tested in the mouse. As shown in Figure 1, a rigorous characterization of color sensitivity requires measurements at multiple SFs. For example, if a neuron has a RF like “single-opponent-A”, it will be more sensitive to S-R (color) than S+R (luminance) contrast, but only at the lowest SFs. In contrast, a “non-opponent” neuron will be most sensitive to S+R at all SFs. We asked whether mouse V1 neurons respond more strongly to S-R or S+R contrast, at each SF in the stimulus set.

To first summarize the raw data, statistics of ΔF/F are given at each SF, for both S-R and S+R contrast (Fig. 3a). Both contrasts elicit a V1 response at each SF, but these statistics begin to reveal a general theme of Figure 3, which is that mouse V1 prefers S-R contrast at the lowest SF, and S+R contrast at higher SFs. Results of an unpaired t-test are inset within the figure at each SF.

**Figure 3:**
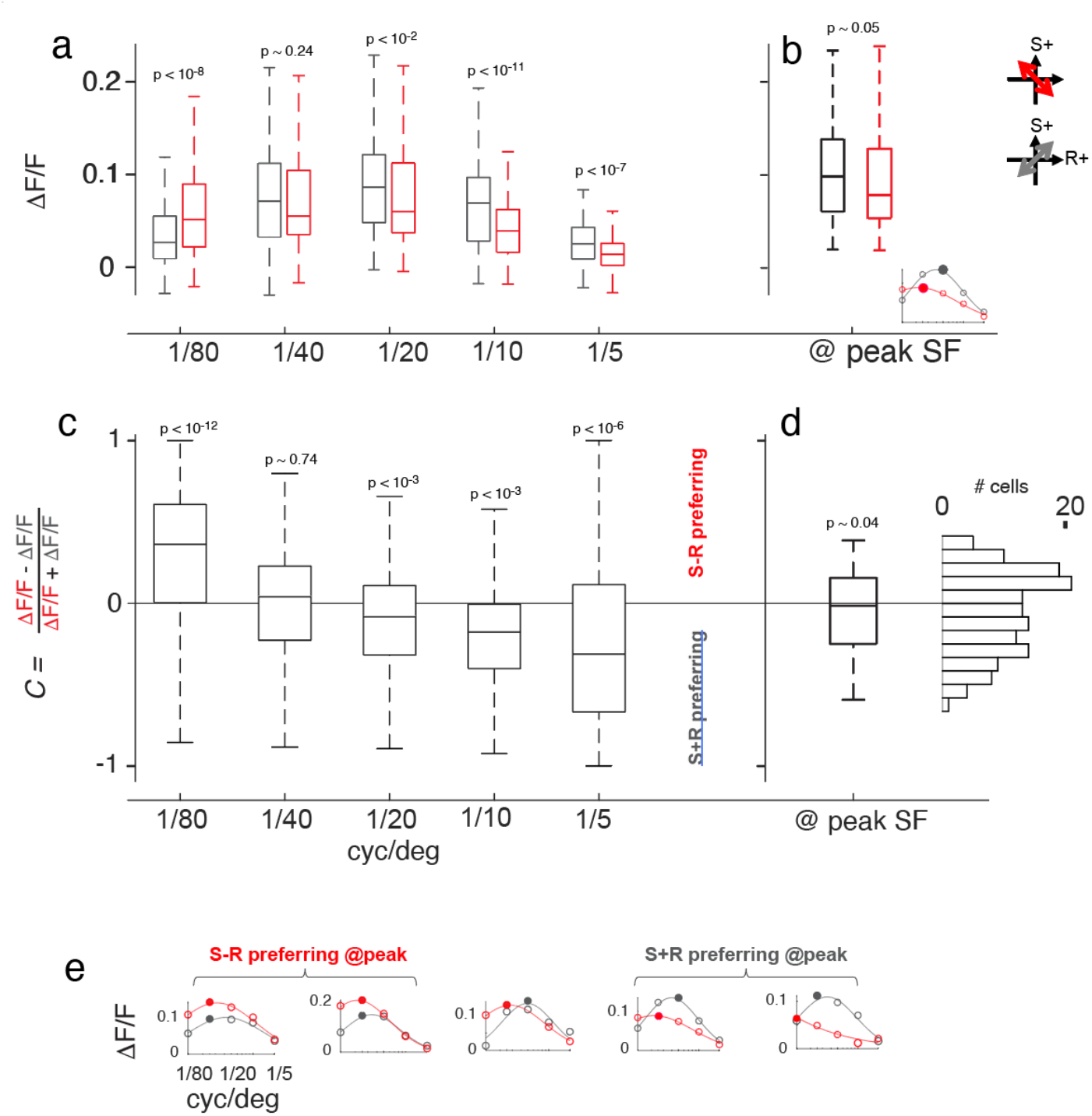
V1’s sensitivity to color and luminance contrast, at each SF. (a) Population statistics of responsiveness to each contrast. For each neuron, ΔF/F at each SF and contrast was calculated as the average over all presented orientations and repeats. The plot shows statistics of ΔF/F for each contrast (S-R in red, S+R in gray), at each of the presented SFs (x-axis). The three horizontal lines of the box plots are quartiles (*q*_*1*_, *q*_*2*_, *q*_*3*_), and the whiskers are located at the the last data point before the “outliers”. Top outliers are greater than *q*_*3*_ + 1.5(*q*_*3*_-*q*_*1*_). Bottom outliers are less than *q*_*1*_ - 1.5(*q*_*3*_ - *q*_*1*_). The p-value at each SF is from an unpaired t-test between the two ΔF/F distributions. (b) Statistics of ΔF/F at the peak SF, for the two contrasts. Inset shows SF tuning from an example neuron, with its two peaks as filled-in data points. The solid line is a Gaussian fit, which was not used for the analysis in this figure. (c) Population statistics of relative responsiveness to S-R and S+R contrast. For each neuron, we calculated preference for S-R over S+R contrast, ‘*C*’ (y-axis), at each SF (x-axis). Population statistics are shown. The p-value at each SF is from a t-test on *C*, identifying whether the mean is different from zero. (d) Population statistics of *C*_*peak*_, where each data point is taken from the peak of each SF tuning curve, S-R and S+R. To the right of the box and whisker plot is a histogram from the same distribution. (e) Six examples of SF tuning and the Gaussian fit, for the S-R and S+R contrast. They are calculated by taking the mean over orientations and repeats.

Next, we calculated each neuron’s preference for S-R over S+R contrast at each SF, ‘*C*’, defined as *C =* (ΔF/F_S-R_ - ΔF/F_S+R_)/(ΔF/F_S-R_ + ΔF/F_S+R_) (Fig. 3c, y-axis). Therefore, *C* = −1 and 1 indicates exclusive response to S+R and S-R, respectively. The distribution of *C* gradually shifts to luminance-preferring values with increasing SF. The mean of *C* at the lowest SF (1/80 c/deg) is significantly greater than 0 (paired t-test; p < 10^−12^). At higher SFs, the mean of *C* is significantly less than 0 (see Fig. 3c for p-values). In summary, mouse V1 prefers color contrast at the lowest SF, and luminance contrast at higher SFs, which likely reflects a meaningful proportion of single-opponent neurons in the population. As will be described, this does not rule out double-opponent or non-opponent neurons, nor necessitate modularity of color and form processing as proposed in the primate ^23^.

To allow simple yet meaningful comparisons in subsequent figures, we reduced each neuron’s preference for color to a single metric, *C*_*peak*_ (Fig. 3d). For a given neuron, *C*_*peak*_ compares the peak response of the S-R SF tuning curve against the peak response of the S+R SF tuning curve; i.e. this entails comparing the response between different SFs if the peak locations differ. The mean and SD of *C*_*peak*_ are 0.04 and 0.25. The distribution looks potentially bimodal, implying that the V1 population can be classified into two groups based on *C*_*peak*_. However, it was not significantly bimodal based on a Hartigan’s dip test (p = 0.84). To compute the p-value, we calculated a Hartigan’s dip statistic ^24^ for *C*_*peak*_, and compared it to the dip statistics from 10K iterations (n = 140 samples; same as Fig. 3d) of a uniform distribution ^25^. The distribution of *C* at each of the 5 SFs (Fig. 3c) also had p-values near one.

We nonetheless segregated the population into “S+R preferring” (*C*_*peak*_ < 0, n = 72) and “S-R preferring” (*C*_*peak*_ > 0, n = 68). The rationale for this segregation is its utility in identifying which of the four opponency models from Figure 1 best describes each neuron. For example, *C*_peak_ is greater than zero for single-opponent-A neurons, but *C*_peak_ < 0 for single-opponent-B neurons. Also, non-opponent neurons have *C*_peak_ < 0, whereas double-opponent neurons have *C*_peak_ > 0.

### The shape of SF tuning depends on opsin contrast

Next, we investigated dependencies of SF tuning parameters on the axis of opsin contrast. The SF tuning curve of each neuron was first calculated as the mean over orientations and repeats, for a given opsin contrast. Next, a Gaussian was fit to each tuning curve. Only neurons where all four of its tuning curves could be fit with a Gaussian (>70% variance accounted) were included in this analysis (n = 140), which is the same population used in Figure 3. To quantify the shape of SF tuning, we began with a metric we refer to as bandpass factor (“BPF”) (Figure 4a-h). BPF quantifies the degree to which SF tuning is lowpass vs. bandpass, based on the Gaussian fit. If BPF = 0, then the Gaussian fit has a peak at the lowest SF (i.e. lowpass). If BPF = 1, the fit is zero at the lowest SF (i.e. bandpass). BPF is particularly useful for classifying each neuron as one of the four opponency models in Figure 1. In addition to BPF, SF tuning was quantified using the peak location of the Gaussian fit (cyc/deg) for each opsin contrast (Figure 4e-h).

**Figure 4:**
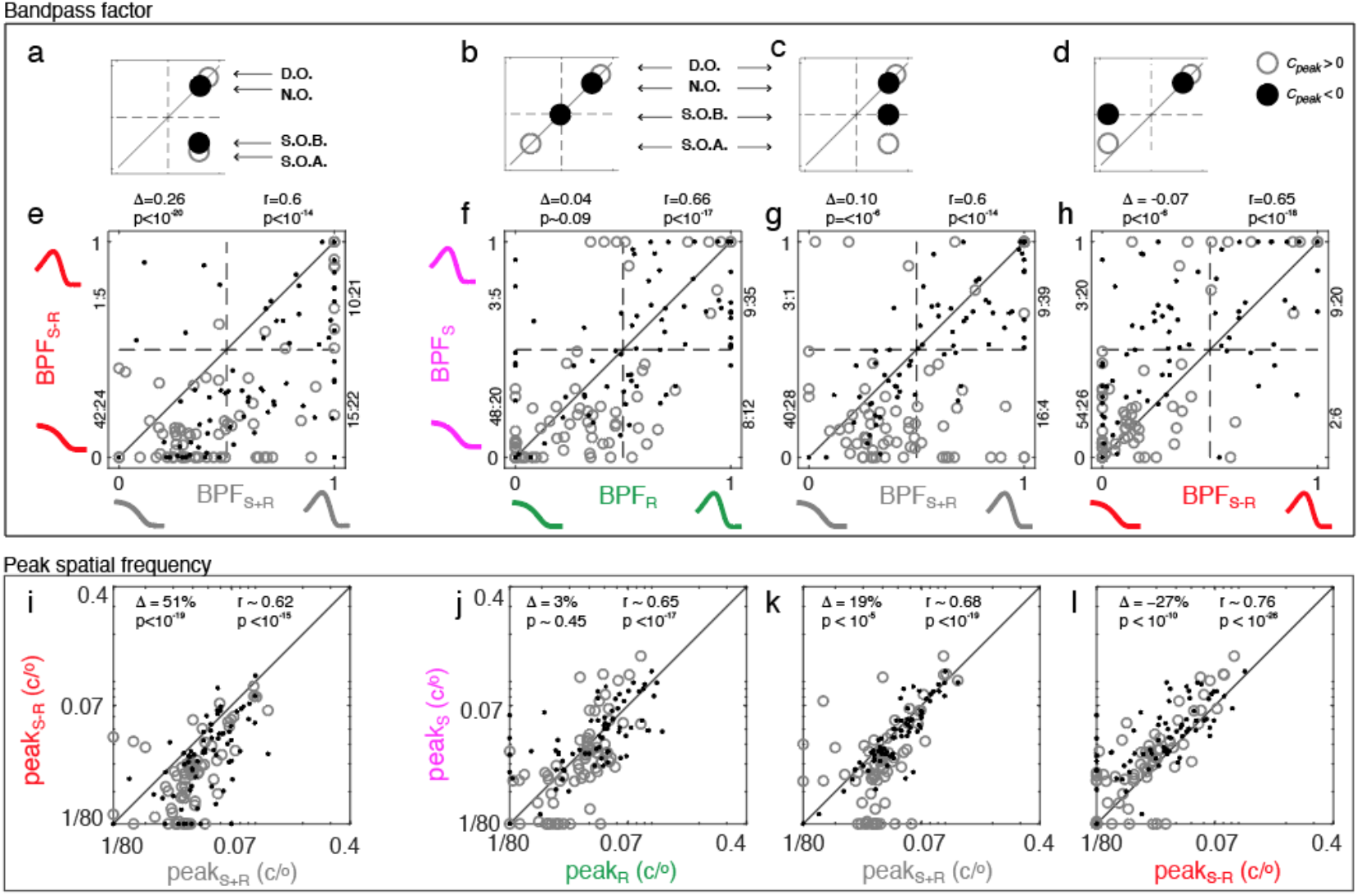
Dependencies SF tuning on opsin contrast. (a-d) These small panels provide an approximate expectation of the four opponency models described in Figure 1. The axes in each panel match the scatter plot immediately below in ‘e-f’, which compare bandpass factor (BPF) between opsin contrasts. Each circle or disk corresponds to an opponency model from Fig. 1: ‘D.O.’ = double-opponent, ‘N.O.’ = non-opponent, ‘S.O.B.’ = single-opponent-B, ‘S.O.A.’ = single-opponent-A. Open circles are color-preferring and closed circles are luminance-preferring. (e) Each data point shows a neuron’s bandpass factor for color (BPF_S-R_) and luminance (BPF_S+R_) taken from the Gaussian fits. The methods for data inclusion (n=140) were outlined in Fig. 2. Unity line is the diagonal. On the top-left,’Δ’ is the mean of BPF_S+R_ −BPF_S-R_, and ‘p’ is from a t-test on Δ. On top-right is the Pearson correlation (‘*r*’) and p-value. To the right or left of each quadrant is the ratio of neurons with open (*C*_*peak*_ > 0) and closed (*C*_*peak*_ < 0) data points. (f) BPF_S_ vs. BPF_R_. Same statistics as in ‘e’ are given around the perimeter. (g) BPF_S_ vs. BPF_S+R_. (h) BPF_S_ vs. BPF_S-R_. (i-l) Same data and layout as in ‘a-b’, but the tuning parameter is the peak location of the fitted Gaussian (cyc/deg). (i) Peak SF for color (peak_S-R_) vs. luminance (peak_S+R_). Unity line is the diagonal. On the top-left, ‘Δ’ is how much bigger peak_S-R_ is than peak_S+R_, as a %. The p-value is from a t-test on log(peak_S-R_)-log(peak_S+R_). On the top-right is the Pearson correlation and p-value comparing log(peak_S+R_) and log(peak_S-R_). (j) Same as ‘e’, but peak_S_ vs. peak_R_. (k) peak_S_ vs. peak_S+R_. (l) peak_S_ vs. peak_S-R_.

The scatter plot in Figure 4e gives each neuron’s BPF for color (BPF_S-R_) and luminance (BPF_S+R_) contrast. The data points sit below the unity line (p < 10^−20^; paired t-test), showing that SF tuning is more bandpass for luminance than color. The smaller panel above (Fig. 4a) gives the predicted location of each of the four opponency models described in Figure 1. Like the data, the opponency models lie between the unity line and the x-axis. Next, Figure 4f shows the BPF of the two opsin-isolating contrasts, BPF_S_ and BPF_R_. They are significantly correlated (p < 10^−17^), and not significantly biased toward either side of the unity line (p ~ 0.09). Consistently, the four opponency models sit on the unity line within these axes (i.e. BPF_R_ ~ BPF_S_) (Fig. 4b). Next, to summarize Figures 4g,h, BPF of the opsin-isolating contrast sits “between” that of color and luminance; i.e. BPF_S-R_ < BPF_S_ < BPF_S+R_, where each of the two inequalities was statistically significant. Again, this dependence of BPF on opsin contrast is consistent with the general layout of the opponency models in the top panels (Fig. 4c,d).

If BPF is replaced with peak SF, the results described above are directionally the same. For one, the peak SF from luminance contrast (peak_S+R_) is greater than the peak SF from color contrast (peak_S-R_) by 51% (p < 10^−19^; paired t-test) (Fig. 4i). Also, peak_R_ ~ peak_S_ (Fig. 4j) and peak_S-R_ < peak_S_ < peak_S+R_ (Fig. 4k,l). It should be noted that a very similar result to peak_R_ ~ peak_S_ (Fig. 4j) was reported in our recent study ^18^.

Lastly, we highlight two important biases in the preference for color over luminance (*C*_*peak*_) within specific quadrants of the BPF scatter plots. First, a sparsity of color-preferring neurons (open circles) in each of the upper-right quadrants of Figures 4e-h provides evidence that few neurons in mouse V1 are double-opponent. Specifically, the modeling panels (Fig. 4a-d) show that color-preferring neurons in the upper-right quadrants of Figure 4e-h are putatively double-opponent, yet 8 neurons (~6%) satisfy this constraint in all 4 axes. At most, 10 neurons (7%) are putatively double-opponent based on Figure 4e alone. Second, color-preferring neurons are more concentrated in the bottom quadrants of Figures 4f-h, consistent with the layout of opponency models in Figures 4b-d. That is, neurons which are lowpass for S-opsin contrast tend to prefer color, which are putatively ‘single-opponent-A’ based on the layout of the opponency models. To assess statistical significance of the relationship between *C*_*peak*_ and SF tuning, we defer to Figure 5, which treats *C*_*peak*_ as a continuous parameter. Specifically, we test the prediction that *C*_*peak*_ is anticorrelated with BPF_S_ and BPF_S_. The results of this test, along with all of the aforementioned trends seen in Figure 4, are later replicated by our random wiring model in Figures 6&7.

**Figure 5:**
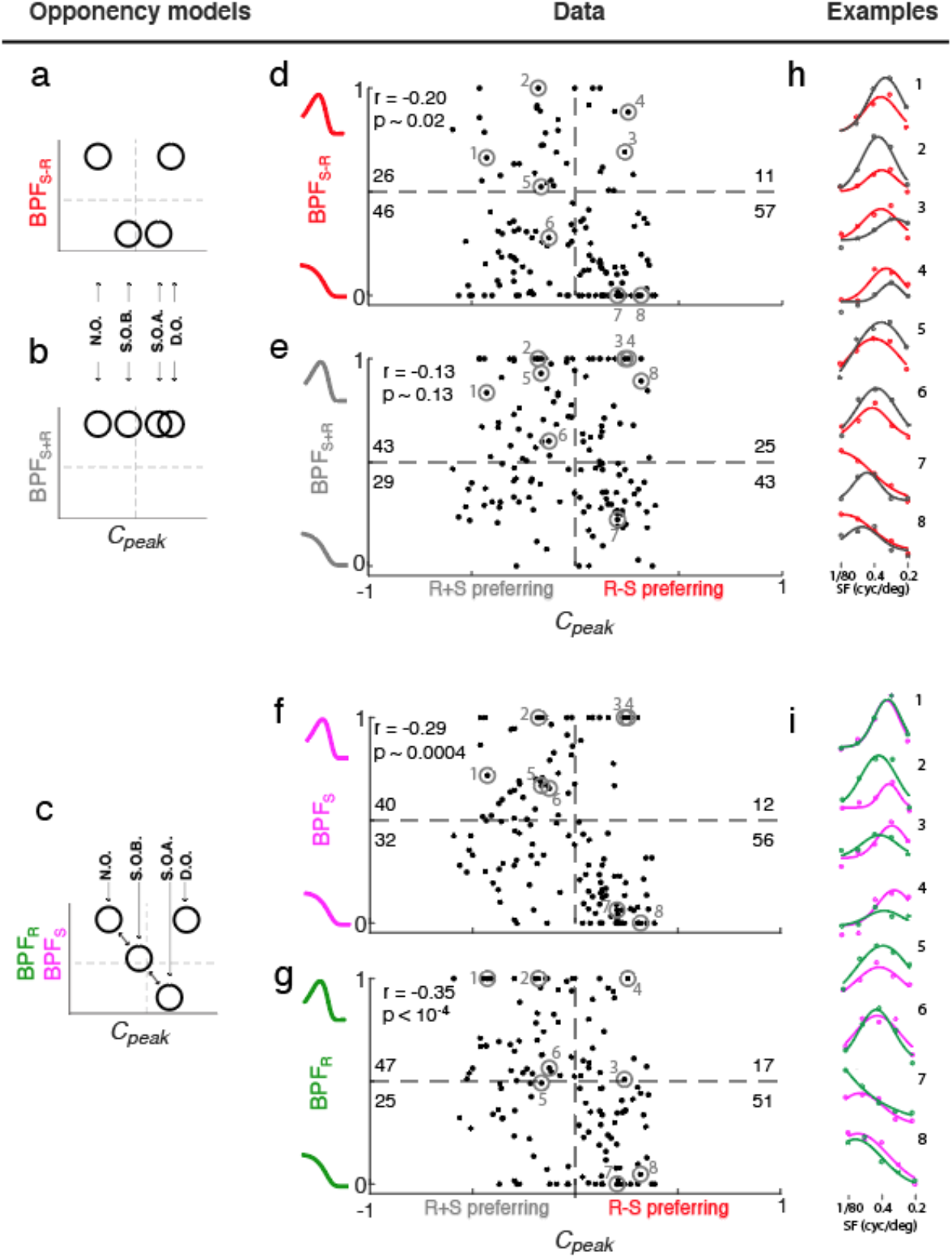
Dependencies SF tuning on color sensitivity. (a-c) These smaller panels illustrate the approximate location of data points from the four opponency models described in Figure 1. ‘D.O.’ = double-opponent, ‘N.O.’ = non-opponent, ‘S.O.B.’ = single-opponent-B, ‘S.O.A.’ = single-opponent-A. The axes are the same as the adjacent scatter plots in ‘d-g’. (d-g) Each panel is from the same population (n=140) as in Figures 3,4, now comparing SF bandpass factor (BPF) of a specified opsin contrast (y-axes) to the preference for color over luminance (*C*_*peak*_) (x-axis). (d) Each data point is a neuron’s BPF for color (BPF_S-R_) and *C*_peak_. At top-left is the Pearson correlation (‘r’) and p-value. Inset within each quadrant is the number of neurons. (e) BPF for luminance (BPF_S+R_) vs. *C*_peak_. Same statistics as ‘d’ are given. (f) BPF_S_ vs. *C*_peak_. (g) BPF_R_ vs. *C*_peak_. (h) Example SF tuning curves and Gaussian fits for S-R (red) and S+R (gray) contrast, which are numbered and circled in each scatter, ‘d-g’. (i) SF tuning curves and Gaussian fits from the same neurons as in ‘h’, but for S (violet) and R (green) contrast.

**Figure 6:**
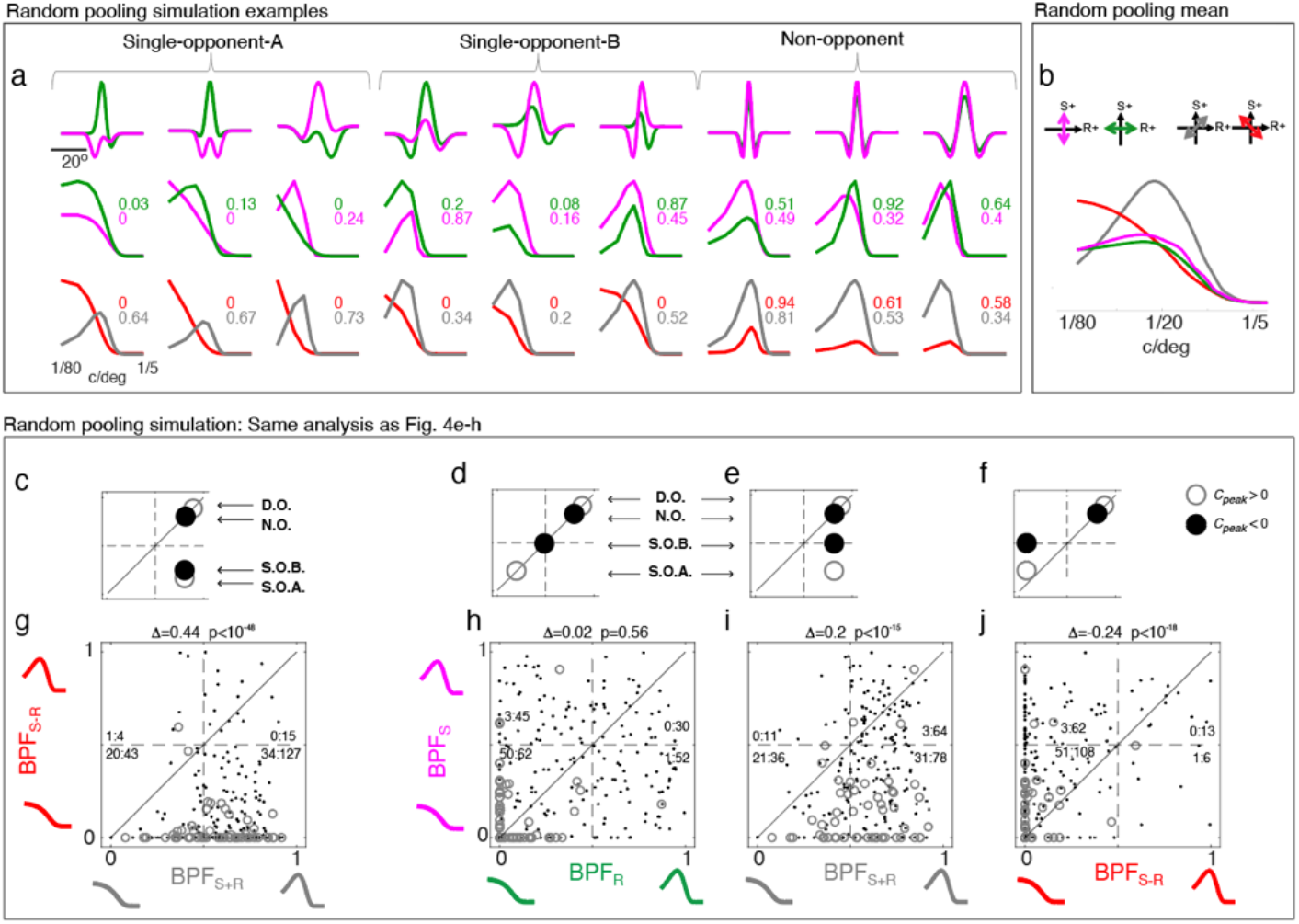
Simulation of results in Fig. 4 with a random pooling model. (a) Each column shows the results of analyzing one simulated neuron that was generated by the random pooling model. The model can generate cells from three opponency classes: single-opponent-A (left 3), single-opponent-B (middle 3), and non-opponent (right 3) cells. The top panels show the spatial S-an R-opsin RF, where the model’s random variables – spatial scale, and the ratio of S-to R-opsin input to each of the three Gaussian subfields – can be more explicitly visualized. The second row shows the Fourier magnitudes of S and R-opsin RFs, to simulate SF tuning. Similarly, the third row is the Fourier magnitudes of S+R and S-R. Inset are the BPFs. (b) The mean SF tuning for each axis in our opsin contrast space, taken from 244 randomly drawn V1 neurons. (c-f) Smaller panels along the top illustrate approximate expectations of the four opponency models described in Figure 1. The axes in each panel match the scatter plot immediately below in ‘e-f’, which compare bandpass factor (BPF) of different opsin contrasts. Each circle or disk corresponds to an opponency model: ‘D.O.’ = double-opponent, ‘N.O.’ = non-opponent, ‘S.O.B.’ = single-opponent-B, ‘S.O.A.’ = single-opponent-A. Open circles are color-preferring and closed circles are luminance-preferring. (g-j) Analysis of 244 simulated neurons, using the same procedures that generated Fig. 4e-h. The titles give the mean difference between BPF on the x-axis and y-axis, and p-value of a paired t-test. Inset within each quadrant is the ratio of color-preferring (open circles), to luminance-referring (closed circles).

### Dependencies of SF tuning on color sensitivity

In Figure 4, the preference for color over luminance (*C*_*peak*_) was characterized as a discrete quantity. Here, *C*_*peak*_ is instead treated as a continuous variable (Fig. 5a-g, x-axes) to examine its joint statistics with SF tuning.

First, we compared BPF_S-R_ to *C*_*peak*_. This comparison is unique in that each opponency model resides within its own separate quadrant (Fig. 5a). There is a weak negative correlation (Fig. 5d), which seems partly due to the scarce sampling in the top-right quadrant, which is the approximate domain of double-opponent cells. The majority of cells are in the bottom two quadrants, which are putatively single-opponent-A and single-opponent-B. Tuning curves and fits are shown for 8 neurons distributed throughout all four quadrants (Fig. 5h), thus exemplifying each opponency model from Figure 1.

Next, we compared BPF_S+R_ to *C*_*peak*_. The opponency models all cluster near BPF_S+R_ = 1.0, irrespective of *C*_*peak*_ (Fig. 5b), which is qualitatively consistent with the data (Fig. 5e) and the insignificant correlation between these parameters. However, there is a substantial proportion of neurons with BPF_S+R_ < 0.5. This can be explained by the idealization of perfectly balanced ON and OFF inputs in the opponency models, which will not be accurate for every neuron. Specifically, the opponency models, as plotted, assume that each neuron always receives a balanced set of ON and OFF inputs, which leads to consistent bandpass tuning for S+R contrast. When the ON or OFF inputs are stronger, the model would predict tuning that is more lowpass for S+R contrast.

Next, we compared the BPF from opsin-isolating contrasts (BPF_S_ and BPF_R_) to *C*_*peak*_. Three of the opponency models (all but double-opponent) sit along a diagonal line (Fig. 5c). Therefore, since the population of double-opponent neurons is relatively small, BPF_S_ and BPF_R_ should be anti-correlated with *C*_*peak*_. This predicted anti-correlation is consistent with the data for both BPF_S_ and BPF_R_ (Fig. 5f,g; see inset statistics).

Finally, Figure 5c is used to illustrate the motivation for developing the random wiring model in the next section (Figs. 6,7). The single-opponent-A neurons are at one extreme, where the ON and OFF channels pool exclusively from either S- or R-opsin in the retina, which yields lowpass SF tuning for S and R contrast, and a preference for color. At the opposite extreme, ON and OFF subfields pool R- and S-opsin in equal proportion to create non-opponent cells, which yields bandpass tuning for S and R contrast, and a preference for luminance. Single-opponent-B simply arise from being somewhere between these two extremes in terms of their R- and S-opsin input ratios. Therefore, by simulating ON and OFF channels that pool at random from the photoreceptors, one can potentially re-create the observed trends in both Figures 4 and 5.

**Figure 7:**
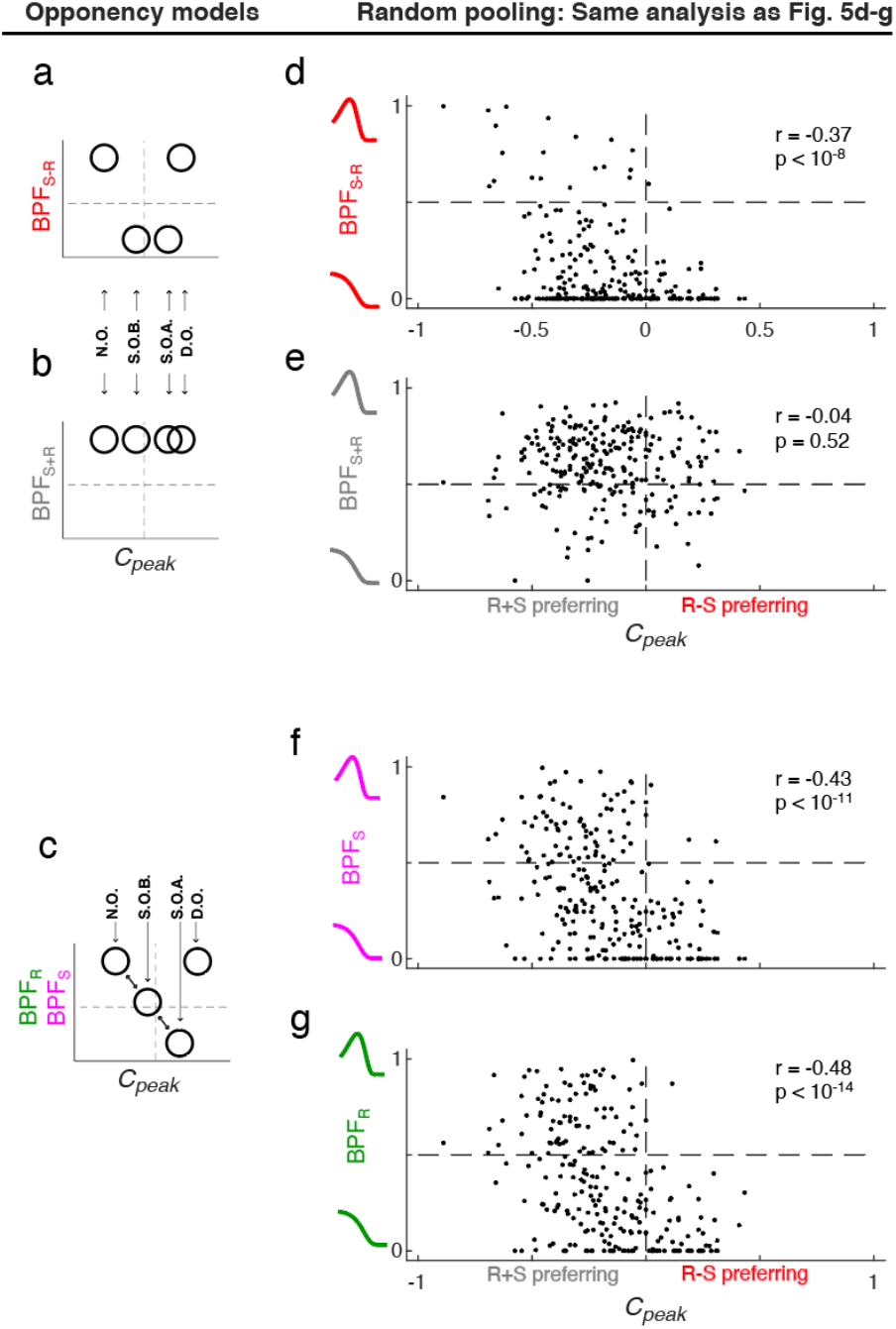
Simulation of results in Figure 5 with a random pooling model. (a-c) Left panels illustrate the approximate layout of opponency models described in Figure 1. The axes are the same as the adjacent scatter plots in ‘d-g’, where the y-axis is the bandpass factor (BPF) of a specified color contrast, and the x-axis is *C*_*peak*_. Each circle or disk corresponds to an opponency model: ‘D.O.’ = double-opponent, ‘N.O.’ = non-opponent, ‘S.O.B.’ = single-opponent-B, ‘S.O.A.’ = single-opponent-A. (d-g) Analysis of 244 simulated neurons (same population as in Fig. 6), using the same procedures that generated Fig. 5d-g. At top-right of each panel is the Pearson correlation (‘r’) and p-value.

### A simple random pooling model describes the data

Next, we constructed a model that explains the observed relationships between SF tuning and opsin contrast in Figures 4 and 5. The model can be briefly summarized as follows. Each neuron starts with a 1D achromatic spatial RF consisting of three adjacent subfields: OFF, ON, and OFF. Next, each of the three subfields is composed of a weighted sum of R- and S-opsin input, where the ratio of the linear weights is chosen at random – this is the “random pooling”. Together, this yields two RFs for each neuron - one for R-opsin and one for S-opsin – that have the same three subfields, but with variable amplitudes. Biological inspiration for the model is described in the Discussion.

Nine example neurons from the random pooling model are shown in Figure 6a (top). The chosen examples illustrate the model’s ability to sample from the three opponency models that were most consistently observed in the actual data. 1) The single-opponent-A examples have the strongest S-R opsin opponency, which is due to the exclusivity of opsin input to the ON and OFF components; e.g. ON is exclusively built from S-opsin and OFF is exclusively from R-opsin. 2) Single-opponent-B examples have input from both opsins to the ON and OFF subfields, which creates SF tuning for S- and R-contrast that is more bandpass than single-opponent-A, in addition to lower S-R sensitivity. 3) Non-opponent examples have similar proportions of S and R-opsin input to each subfield, which creates similar SF tuning across all four opsin contrasts. i.e. they are much more separable. Next, we assessed the distributions of these three opponency models from a randomly sampled population.

244 neurons were randomly generated using statistical parameters described in the Methods. Using this simulated population, we repeated the data analyses comparing BPF across opsin contrasts (Fig. 4e-h). For ease of qualitative and quantitative comparison, the model opponency panels in Figure 4a-h are reshown in Figure 6c-j. Overall, the random pooling model does well at capturing key features of the data. To summarize, the average simulated neuron responds to color contrast (S-R) with lowpass SF tuning, to luminance contrast (S+R) with bandpass SF tuning, and to opsin-isolating contrast (S or R) somewhere between (Fig. 6b,g-j). Furthermore, like the actual data, there are clear biases in the location of color-preferring neurons - simulated cells that prefer color (open circles) dominate the bottom two quadrants of Figure 4h-j.

In Figure 7 the same simulated dataset was used to compare BPF to *C*_*peak*_. The layout and analysis procedures mimic Figure 5. Perhaps most importantly, the simulated population clearly captures the data’s negative correlation between BPF of opsin-isolating contrast and *C*_*peak*_ (Fig. 7f,g). Furthermore, like the data, there is a weaker correlation between BPF_S-R_ and *C*_*peak*_ (Fig. 7d) and a lack of significant correlation between BPF_S+R_ and *C*_*peak*_ (Fig. 7e)

## Discussion

V1 is much more sharply tuned for spatial form than its subcortical inputs, and an important question is how this spatial transformation becomes jointly encoded with color. Building on recent work showing rod-cone opponency in the ventral portion of the mouse retina ^1–3^, we investigated the joint representation of color and spatial form in the upper visual field representation of mouse V1. On average, we find that V1 neurons in the upper visual field are tuned to lower SFs of color contrast than luminance contrast, which is consistent with a model of single-opponency between rods and cones, as classically defined ^12^; i.e. “single-opponency-A” (Fig. 1). A single-opponent-A neuron has ON and OFF channels that pool exclusively from one of the two photoreceptors, making it lowpass for opsin-isolating stimuli and preferring color (e.g. Fig. 4f, bottom-left quadrant). Neurons fitting this classic single-opponency criteria are about 34% of the population, but our data suggest that they are not the result of selective pooling by ON and OFF channels. Rather, they are more parsimoniously described as one side of a continuum of V1 neurons, which arise from random draws of an un-selective pooling mechanism. At the other side of this continuum, the ON and OFF channels each pool balanced quantities of the two opsins, which yield non-opponent cells (e.g. Fig 4f, upper-right quadrant, closed circles; 25%).

By implementing a random pooling model, we were able to account for other trends in the data as well. For example, the model recapitulates the inseparability of SF tuning across opsin contrast – SF tuning is “lowest” for color, highest for luminance, and somewhere between for opsin-isolating contrast (Fig. 4e-h vs. Fig. 6g-j). Furthermore, the model predicts the negative correlation between color sensitivity and BPF in response to opsin-isolating gratings (Fig. 5f,g vs. Fig. 7f,g). Finally, the random pooling model is consistent with more recent recordings from subclasses of retinal ganglion cells in the mouse and primate ^1,17,26^.

### Why UV-green color opponency in the mouse?

As the sun sets, irradiance at all UV-to-IR wavelengths is heavily attenuated, but longer wavelengths are attenuated the most. What remains until sunrise is a band of spectral irradiance between roughly 300 and 550 nm ^27^. The sensitivity of photoreceptors in the rodent retina covers this range quite well, thus suiting their nocturnal habits. But why would UV-visible color vision be specialized for the upper visual field? It may be to distinguish monochromatic UV contrast from chromatic UV-green contrast. UV radiation alone is heavily scattered by the atmosphere and readily absorbed by most surfaces, which allows for “silhouette detection” in the upper visual field ^5^. But without rods to “paint” less threatening objects green, little may be discerned in a more cluttered upper field. In other words, uniquely monochromatic input may be the most threatening signal, whereas color is less threatening. These hypotheses warrant future behavior experiments that not only vary parameters of form and motion via rod input ^6^, but also color.

As for the lower visual field, since the light reaching the eye has been reflected, UV input is relatively weak. The lower visual field nonetheless contains a sparse sampling of pure S-cones, whose input may be compared to that of M-opsin and rods for efficient foraging and identification of marked territory ^2^.

What about disadvantages of UV-green sensitivity? One example is potential damage to the retina from UV. Naturally, transmittance of UV in the rodent eye is high. However, being nocturnal and having a relatively short life, any accumulated UV damage to the mouse retina is likely minimal compared to that in a larger diurnal animal ^28^. Another potential problem is chromatic aberration, which will place the UV and green contributions of an image at different focal planes. Chromatic aberration is particularly strong at these shorter wavelengths, forcing a substantial fraction of the light spectrum to be out of focus at the retina. However, there are a few things to consider. First, UV-visible chromatic aberration in the mouse eye may not be a limiting factor in a visual system with such poor resolution. For example, if 550 nm is optimally focused at the mouse retina, this creates approximately 20 diopters of aberration at 360 nm ^29^ (see their Fig. 5). Twenty diopters is rather large, as it roughly equates to a “blur function” diameter of 1° of visual field ^30^ (see their Eq. 1), giving a cutoff SF of around 0.5 cyc/deg. However, mouse V1 has very little sensitivity to SFs out to 0.5 cyc/deg. Therefore, chromatic aberration in the mouse eye is likely to occur at SFs that are ultimately filtered out by later stages of the early visual system. A separate study in the rat using numerical and experimental methods showed similar results as our approximate calculations above for the mouse - the maximum aberration-induced blur in the rat eye is about 60 microns, or ~1° of visual field ^31^, which is around the upper limits of resolution at the level of rat V1 ^32^. Regarding any residual effects of chromatic aberration that reach cortex, it’s also worth noting that the distance to objects in the mouse’s upper visual field, when above ground, are effectively at “infinity”. This keeps the aberration of a given object invariant to changes in its proximity, which limits potential confounds for downstream processing. In summary, the pros seem to largely outweigh any cons of a UV-visible color system in the rodent.

### Measuring opponency from the phase of response

We did not report the phase of neural responses, only the mean amplitude. However, a common strategy for measuring color opponency is to compare the response phase at the temporal frequency, between gratings of different opsin contrast. Given the population’s preference for color (S-R) at 1/80 cyc/deg (Fig. 3), we predict that the response to this SF has a phase differential of 180° between S-opsin and R-opsin gratings. Unfortunately, a combination of factors made a rigorous analysis of phase opponency overly challenging. For one, most F1-amplitudes at the stimulus frequency were weak, possibly due to the cells being more “complex”, along with the relatively slow time course of the calcium signal. Furthermore, the PMT in our two-photon microscope was detecting subtle leakage of the green LED in the visual stimulus, which has a 180° phase shift between the R- and S-opsin gratings. Ultimately, this created an artifactual phase opponency that directly aligns with the aforementioned prediction. Importantly, the amplitude of this artifactual modulation does not affect the mean used in the rest of the analyses.

### Orientation tuning selectivity vs. opsin contrast

Orientation tuning becomes more selective in V1 and its joint representation with color is also relevant to the general problem of color and form integration. Here, we designed our experiments to assess color and SF tuning. SF tuning gives more direct interpretations of the data with respect to the opsin-opponency models (Figure 1), and thus the underlying mechanisms of color tuning in V1. Specifically, only four orientations were presented (Δ = 45°), which kept each experiment to a reasonable length, while also ensuring that most neurons were well driven by at least one orientation. We believe that the study is underpowered for a thorough characterization of the joint representation of color and orientation selectivity. Specifically, the sample spacing (45°) is too large to capture systematic changes in the orientation half-bandwidth of many neurons in mouse V1.

We can nonetheless offer predictions about orientation selectivity in the context of the Figure 1 models. In summary, orientation bandwidth is predicted to be correlated with SF tuning - lower BPF implies broader orientation tuning. For example, single-opponent neurons are predicted to be lowpass in SF and have broader orientation tuning, for color contrast. Similarly, double-opponent neurons and non-opponent neurons are predicted to have sharper orientation tuning along any axis of opsin contrast. More generally, in the polar 2D Fourier plane (SF vs. orientation) with SF = 0 at the origin, a trivial consequence of lowpass SF tuning is broader orientation tuning ^10^.

### Orientation tuning preference vs. opsin contrast

The mouse retina has a gradient of cone-opsin expression along its dorsoventral axis, with M-and S-opsin dominating the dorsal and ventral retina, respectively ^33^. Recordings from retinal ganglion cells show that when center-surround RFs pool from this cone-opsin gradient, unselectively, color opponency arises ^34^. What about V1 RFs that have oriented and Gabor-like RFs? Like retinal ganglion cells, V1’s opponency from the cone-opsin gradient will be highest in the transition zone (i.e. near the horizontal midline). In addition, a V1 neuron’s color opponency from this mechanism will depend on its orientation preference. For example, a Gabor-like RF that is selective for vertical edges will not exhibit opponency from co-expressing cones, regardless of its retinotopic position (Supp. Fig. 1b). On the other hand, when a Gabor located near the midline is selective for horizontal edges, we expect cone-opsin opponency to arise since the ON and OFF subfields will pool unique ratios of M and S opsin (Supp. Fig. 1a). The spatial RF due to this mechanism of opponency will look like ‘single-opponency-B’ (Fig. 1).

We tested for opponency from the cone opsin gradient in our data. We compared a metric for inseparability of color and form - the distance of a data point in Figure 4i from the unity line - against preferred orientation. In other words, the inseparability metric tells if the neuron is tuned to higher SFs for luminance than color. This metric provides a convenient summary of opponency because it is higher for single-opponent neurons. The scatter plot is shown in Supplemental Figure 1c. Interestingly, this yielded the expected trend from the cone-opsin gradient (r = −0.25; p = 0.002); neurons tuned closer to horizontal were more inseparable than neurons tuned closer to vertical. However, the effect is weak and even neurons tuned to vertical are inseparable for color and form, implying that most of the inseparability is from rod and cone integration, not the cone-opsin gradient. See Supplemental for additional detail.

### Biological validity of the random pooling model

A more biologically-inspired explanation of the random pooling model is as follows. Each V1 neuron receives input from ON and OFF lateral geniculate cells, which creates a multimodal spatial RF (and thus bandpass SF tuning curve) for luminance contrast ^9,11^. Moving further upstream, the ON and OFF channels each pool random amounts of S- and R-opsin in the retina. This implies that S- and R-opsin contributions to a RF will have the same polarity within a given subfield (i.e. both ON, or both OFF), as shown in the majority of retinal ganglion cells in the ventral retina ^1^. The integrated amount of S-opsin by the ON channel creates its sensitivity (i.e. ON subfield amplitude) to S contrast, and similarly for R-opsin and R contrast. The OFF subfield is generated independently of the ON subfield, but uses the same statistics to pool from R- and S- opsin.

Unlike classic depictions of single-opponent V1 neurons ^12^, our model does not require that ON and OFF subfields systematically pool from the photoreceptor mosaic, which seems consistent with more recent studies in the retina ^1,17,26^. We also note that results of the model’s output are quite robust to an overall bias in S or R sensitivity. For example, when the simulation is re-run with all the R-opsin RFs scaled down by a factor of 2x, such as during brighter states of adaptation, the results remain similar to those in Figures 6 and 7. The difference is a slight increase in color-form separability; e.g. data points in Figure 6g (BPF_R-S_ vs. BPF_R+S_) move toward the unity line. Ultimately, once rods are fully saturated, opponency is gone and the tuning becomes completely separable. Below, we elaborate on the issue of how graded rod saturation affects our data and the separability of color and form.

### Graded discriminability between color and luminance contrast in the mesopic regime

The degree to which a mouse V1 neuron is single-opponent – and thus inseparable for SF and color – is in flux with the balance of rod and cone input induced by a given level of light adaptation. Below, we explain how a graded reduction in the balance of rod and cone inputs leads to reduced discriminability between color and luminance contrast by the V1 population. Figure 1 is used as a guide, and new statistics from our data are provided.

As light levels fall to the lower end of the mesopic range, R-opsin sensitivity will dominate over S-opsin sensitivity. With this imbalance, the SF tuning for S-R and S+R contrast will begin to match the SF tuning for R-opsin contrast. The results are analogous for an increase in background intensity, where the S-opsin dominates. In other words, the prediction is that the RF becomes more separable in color and form with greater opsin imbalance. Consistent with this, we found more separable tuning when we limited analyses to neurons with imbalanced (1/3 > %R > 2/3) instead of balanced (1/3 < %R < 2/3) S and R input. To be clear, the balanced population was used for Figures 3-5. When reanalyzed with the imbalanced population, tuning was still inseparable, but less so. For instance, the change in peak SF between S-R and S+R contrast was 29% (compare to 51% in Figure 4i).

Under complete photopic or scotopic adaptation, a single opsin encodes the upper visual field, which cannot discriminate S-R and S+R contrast because separable color and spatial tuning of each neuron leads to the same pattern of population activity. Conversely, in the mesopic regime, two images with the same form but different color will generate a different pattern of population activity because the RFs are inseparable in color and form. Such differences in the cortical population response could mediate perceptual discrimination between color and luminance contrast ^4^.

### Generalizing results to complex cells

The first three opponency models in Figure 1 are constrained by flanking ON and OFF subfields and are thus consistent with the classic depiction of a “simple cell”. The fourth model, double-opponency, is also a simple cell in the sense that each subfield encodes one direction of color space. All four of the opponency models in Figure 1 are simple cells based on this broad latter definition. However, many cells in mouse V1 are complex ^11^, so it is important to address how our characterizations of the data - based on the models in Figure 1 - might generalize to complex cells.

Each opponency model in Figure 1 has a complex cell counterpart, and the “energy model” ^35^ provides a useful guide for understanding its functionality. In its most basic form, the energy model consists of two simple cells with unique phase preference (but otherwise similar RF properties), whose outputs are squared and then added to create a complex cell. The chromatic SF tuning curves of this theoretical complex cell will depend heavily on how they are measured. In this study, we used the *mean response* to drifting gratings (as opposed to the *max*, or *F1*), in which case the model complex cell has the same SF tuning as its simple cell inputs, for each axis of R and S contrast space. That is, a complex cell that is “downstream” of any of the four simple cells shown in Figure 1d will produce the same SF tuning curves as those in Figure 1e,f, when mean response is used. For example, two non-opponent simple cells, in phase quadrature, combine to produce a complex cell that has the same SF tuning curves as each of its simple cell inputs. This includes the bandpass SF tuning for each opsin contrast, along with a preference for S+R over S-R contrast. In summary, our characterizations of opsin opponency using mean response are expected to generalize to complex cells.

One can further generalize to a complex cell that pools from two or more *different* opponency models in Figure 1. In these cases, the SF tuning of each opsin contrast – using the response mean - can be modeled as a linear superposition of those from the simple cell inputs. Subtle inconsistencies with this notion are related to the shape of the simple cell output nonlinearities, and not related to the relative phase of the inputs.

But these generalizations to complex cells do not hold when the max or amplitude (e.g. “F1”) of the response are used to measure SF tuning. For instance, the output of two single-opponent neurons that have opposite sign – i.e. one S-R and the other R-S - could be combined to generate a neuron that is more bandpass for color contrast, and thus double-opponent, when response amplitude is used.

### In search of double-opponency and bandpass tuning for chromatic edges

Double-opponency is a hypothesized mechanism by which chromatic edge detection is encoded in the primate visual system ^12,36^. Although most studies seem to agree on the most basic definition of a double-opponent neuron - flanking single-opponency of opposite polarity - the criteria for identifying them differs widely ^12,27,34,35^. Based on our simulations of linear RFs, along with our available measurements, it is sensible to define a double-opponent cell as one that 1) prefers color over luminance contrast (i.e. *C*_*peak*_ > 0), and 2) has bandpass SF tuning for all axes in the plane of S and R contrast space. These criteria are related, but not identical to prior studies, such as in ^13^. Our simple criteria seem to generalize quite well when modeled. For instance, one can change the spatial scale and/or amplitude of either the R or S spatial RF, for the double-opponent model Figure 1d, but the SF tuning curves still obey the two criteria mentioned above – as they should since the RF is still technically double-opponent. Its only when the polarity of the R- and S-opsin RFs are the same (i.e. non-opponent) that *C*_*peak*_ is greater than zero. Of course, classification errors will arise due to noise or deviation from the linear model, particularly for cells with *C*_*peak*_ near zero. Nonetheless, our general conclusion that few cells in mouse V1 are double-opponent is likely to hold.

Additional double-opponent RFs could be built in downstream areas by combining V1’s single-opponent outputs ^37,38^. This could arise through simpler linear-nonlinear integration, like in the energy model described above, but requires that a metric like F1 amplitude be used instead of the mean. If the mean response to drifting gratings is used (as in this study), inhibition is required within a more complicated mechanism like “push-pull” ^39^ or local feedback. Whether response mean or F1 amplitude is the most useful metric depends on the given perceptual task. We believe these issues warrant future theoretical and empirical investigation on the transformations of color and form beyond mouse V1, along with its behavioral correlates.

## Methods

### Animal preparation for surgery and imaging

All experiments were approved by the University of Texas at Austin’s Institutional Animal Care and Use Committee, which maintains AAALAC accreditation. All methods were carried out in accordance with relevant guidelines and regulations. 8 male and 5 female wild-type mice (C57BL/6; aged 2-5 months) were used.

For all surgical procedures, mice were anesthetized with isoflurane (3% induction, 1-1.5% surgery), and given a pre-operative subcutaneous injection of analgesia (5mg/kg Carprofen) and anti-inflammatory agent (Dexamethasone, 1%). Mice were kept warm with a water-regulated pump pad. Each mouse underwent two surgical procedures. The first was the mounting of a metal frame over the visual cortex using dental acrylic, which allowed for mechanical stability during imaging. The second was a craniotomy (4-4.5mm in diameter) over V1 in the right hemisphere, virus injections, and a window implant (3-4mm glass window). Surgical procedures were always performed on a separate day from functional imaging. On a day between the frame implant and virus injections, we identified the outline of the V1 border by measuring the retinotopic map with intrinsic signal imaging (ISI) through the intact bone^40,41^.

The virus (pAAV.Syn.GCaMP6f.WPRE.SV40, Addgene viral prep # 100837-AAV1)^42^ was delivered using a Picospritzer III (Parker) or Nanoliter injector (WPI) to 2-3 sites in V1 with 0.25-0.5ul per site. Once the injections were complete, craniotomies were sealed using Vetbond and dental cement with dual-layered windows made from coverglass (4mm glued to 5mm, Warner Instruments), and covered between imaging sessions.

Common to all functional imaging procedures, isoflurane levels were titrated to maintain a lightly anesthetized state while the mouse laid on a water-circulating heat pad. However, mapping the V1 retinotopy using ISI is typically more tolerant to high isofluorane levels. For ISI, isoflurane levels were set to 0.25-0.8% (typically 0.5%). For two-photon imaging, each mouse was given chlorprothixene (1.25 mg/kg) intramuscularly and isoflurane levels were adjusted to 0.25-0.5%. Silicone oil (12,500 cst) was periodically applied to the stimulated (left) eye to prevent any dryness or damage to the eye. Silicone oil allows optical transmission of near-UV (nUV) and visible wavelengths. Additionally, the unstimulated (right) eye was coated with Vaseline and covered with black tape during imaging.

### Imaging

For ISI, images were captured using a Pco Panda 4.2 camera with Matlab’s image acquisition toolbox, at 10-to-20 frames/sec, and 2×2 binning. An X-Cite 110LED (Excelitas) was used for illumination. A longpass colored glass filter (590nm) was placed over the light guides to illuminate the brain, and a bandpass filter (650 +/− 25 nm) was on the camera lens.

Two-photon calcium imaging was performed with a Neurolabware microscope and Scanbox acquisition software. The scan rate varied between 10-15 frames/sec, scaling with the number of rows in the field-of-view. A Chameleon Ultra laser was set to 920 nm to excite GCaMP6f. A Nikon 16x (0.8NA 3mm WD) or Olympus 10x (0.6NA 3mm WD) objective lens was used for all imaging sessions.

### Visual stimuli

#### Setup overview

Two monochrome LED projectors by Texas Instruments (Keynote Photonics) with spectral peaks at 405 nm (nUV) and 525 nm (green) were used to generate drifting sinewaves with contrast in the S- and R-opsin plane. The refresh rate was 60 Hz. The rear projector screen was made of Teflon, which provided a near-Lambertian surface^18^. One projector was aligned to the other with an affine transformation in Matlab. The transformation was identified by a user’s manual selection of several points on the screen. Stimuli were coded using the Psychophysics Toolbox extension for Matlab^43,44^.

The mouse was positioned so that its left eye was vertically centered on the projector screen, and the perpendicular bisector from the mouse’s eye to the screen was between 8 and 10 cm. Next, the screen was angled at 30° from the mouse’s midline. The final constraint was to align the front edge of the screen to the mouse’s midline. The screen size was 43 cm high x 23.5 cm wide, which gives approximately 135° x 105° of visual angle.

#### Retinotopic mapping stimulus

The same drifting bar stimulus was used to map retinotopy for both ISI and two-photon imaging. This method is an extension of the one used in Kalatsky and Stryker ^21^, and described in detail in Marshel et al.^20^. To summarize, a periodic drifting bar on black background was modulated by a contrast-reversing checkerboard, and shown in four cardinal directions. The bar’s speed and width were dependent on the screen location such that the temporal phase of cortical responses, at the stimulus frequency, could be directly mapped to altitude and azimuth coordinates in a spherical coordinate system. The speed of the bar was ~5°/s and its width was ~12°. The checkerboard squares were 5° wide and reversed contrast at a rate of 2 Hz. Both green and nUV projectors were shown at maximum contrast to produce the checkerboard. Each drift direction was shown at ~0.055 HZ for wide-field imaging and ~0.042 HZ for the 2-photon calcium imaging setup. There were 2-to-3 120 sec trials, for each drift direction.

#### Color-SF stimulus

Full-field drifting sinewave gratings at variable SF, color, and direction were shown. The ensemble contained five spatial frequencies (0.0125, 0.025, 0.05, 0.1, 0.2 cyc/deg), 4 axes of S and R contrast space (S, R, S-R, S+R), and 8 equally-spaced drift directions (i.e. 4 orientations). Five repeats of all combinations were shown, totaling 5×4×8×5 = 800 trials. The experiment was split in half. One half was limited to the S and M axes of color space, and the other half was limited to the S-R and S+R axes of color space. Stimuli were otherwise randomly interleaved. Each grating was shown for 3 sec, followed by 2 sec of a uniform “gray” screen that matched the mean spectral radiance of the gratings.

At least 10 minutes before recording, the pupil was fully dilated with 1% tropicamide, and the retina was adapted to the gray background of the stimuli. The gray background is estimated to yield approximately 200×10^5^ photoisomerizations/rod/sec, calculated below. In a prior study, we found that this rod isomerization rate yields V1 responses that are mediated by both rods and cones ^18^. As shown in Figure 2, data yield was limited to cells that are similarly responsive to S- and R-opsin isolating stimuli.

### Calculation of rod isomerization rate

In a recent study, we described the calculation of the rod isomerization in this same preparation ^18^. We summarize these calculations here. We start with the calculation of spectral irradiance at the retina as follows:

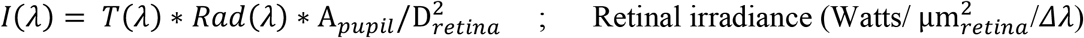

*T*(*λ*) is transmittance through the mouse lens at each wavelength ^45^. *Rad*(*λ*) is the spectral radiance of the gray background, measured with a PR655 spectroradiometer. A_*pupil*_ is the diameter of the fully dilated mouse pupil, and D_*retina*_ is the diameter of the retina ^46^. Next, we convert Joules to quanta as

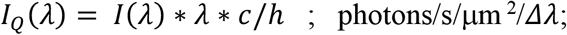

where *c* is the speed of light and *h* is Planck’s constant.

To compute isomerization rate, *R*^∗^/*receptor*/*s*, we take the dot-product between the quantal retinal irradiance, *I*_*Q*_(*λ*), and the absorption spectrum, *a*_*c*_(*λ*), and scale by the sample period of the spectrum *Δλ*:

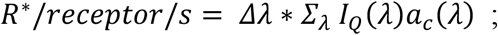

To get *a*_*c*_(*λ*) in μm^2^, we started with the unitless absorption spectra of rods from Govardovskii et al.^47^ and then scaled by the end-on collection area at the peak wavelength which may be approximated as 0.85 μm^2^ for rods ^48^.

### Calibration of cone-opsin contrast

#### S- and M-opsin isolating gratings

Green and nUV LED sinewaves were combined to produce contrast along one of four axes in cone-opsin space (Fig. 1f). As shown below, S-opsin and R-opsin contrast can be described as a function of the rear-projected spectral radiance of the nUV (*R*_*UV*_(*λ*)) and Green (*R*_*G*_(*λ*)) LEDs, the amplitude of their drifting sinewaves (*a*_*UV*_, *a*_*G*_), and the S- and R-opsin sensitivity functions (*h*_*S*_(*λ*) and *h*_*R*_(*λ*)):

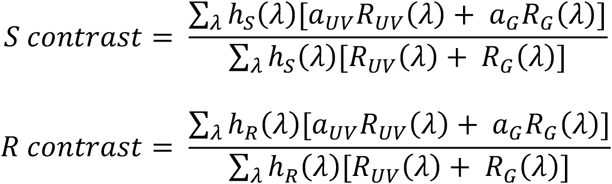

Spectral radiance was measured using a PhotoResearch PR655. Opsin sensitivity functions are taken from Govardovskii et al.^47^ The above equations were used to solve for *a*_*UV*_ and *a*_*G*_, for each of the four color axes in this study: S, R, S-M, and S+M. However, the total opsin contrast along each axis must first be constrained. Total opsin contrast, 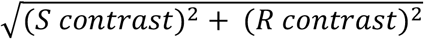, was set at 60% for each axis. For example, along the S-axis of color space, *S contrast* = 60% and *R contrast* = 0%, whereas for the S+R and S-R axes, each opsin had a contrast of 43%. 100% total contrast was not used because it was not achievable in all axes. The formulae above consist of two equations and two unknowns (*a*_*UV*_ and *a*_*G*_), which can be solved for each color axis.

### Rod and cone random pooling model

Each simulated neuron in Figures 6 and 7 starts with a monochromatic RF that is the addition of three 1D Gaussians: G(x) –G(x-2σ)/2 –G(x+2σ)/2. Here, G(x) is the Gaussian as a function of degrees of visual space (x), and has a standard deviation of σ. The first Gaussian is the ON subfield, and the other two are the flanking OFF subfields. The OFF subfields are attenuated by a factor of 2, and shifted to the right and left by 2σ, relative to the ON.

RFs were scale invariant. The peak SF, *f*_o_, of a RF under these constraints is approximately equal to (*πσ*) ^−1^. To simulate the population of spatial RFs, we sampled from a gamma distribution of *f*_o_, which yields the spatial RF of ON and OFF Gaussians according to *σ* = (*πf*_*O*_) ^−1^. The gamma distribution of *f*_o_ was comparable to the data, with a K (i.e. “shape”) of 10, and a “scale” of 0.05/K, which sets the mean to 0.05 cyc/deg.

Next, each monochromatic ON or OFF subfield was parsed into a linear combination of input from R-opsin and S-opsin. Specifically, each subfield sampled from a binomial distribution (n = 3, p = 0.5) to create the relative strength of input from S- and R-opsin. The three independent binomial draws, one for each subfield, gives the Gaussian amplitudes of the three S-opsin subfields, A_S-ON_, A_S-OFF-right,_ A_S-OFF-left_. The Gaussian amplitudes of the three R-opsin subfields are simply n minus the S-opsin amplitudes: A_R-ON_ = n – A_S-ON_, A_R-OFF-right_ = n – A_S-OFF-right_, and A_R-OFF-left_ = n – A_S-OFF-left_. We define the two opsin RFs as “RF_Sopsin_” and “RF_Ropsin_”, where RF_Sopsin_ = G(x) *A_S-ON_ – G(x-2σ)/2 *A_S-OFF-right_ – G(x+2σ)/2 *A_S-OFF-left_.

To simulate SF tuning curves in response to each contrast direction, we took the Fourier transform magnitudes. For instance, the SF tuning of S+R and S-R contrast was taken as the Fourier transform magnitude of RF_Sopsin_ + RF_Ropsin_ and RF_Sopsin_ - RF_Ropsin_, respectively. Next, we quantified SF tuning with identical procedures used on the real data; viz. a Gaussian function was fit, from which the BPF was calculated.

### Modeling V1 RFs that pool from the cone-opsin retinotopic gradient

Supplemental Figure 1a,b show model RFs in order to help interpretate the data in Supplemental Figure 1c. The model RFs start with an achromatic 1D difference-of-Gaussian (DoG). The Fourier transform of the RFs have a peak SF at 0.05 cyc/deg. Next, this achromatic 1D RF is decomposed into separate S-opsin and M-opsin RFs using a model of the opsin gradient given in ^7^. We denote the opsin gradient as %M(*v*), which is the concentration of M-opsin at retinotopic location ‘*v*’. To obtain %M(v) from ^7^, the domain was converted from millimeters on the retina into degrees of visual field (*v*). When the RF is tuned for horizontal (Supp. Fig. 1a), the 1D DoG is a function of *v*, and thus scaled %M(*v*) to get the M-opsin RF, and scaled by 100-%M(*v*) to get the S-opsin RF. The SF tuning curves for each opsin contrast were then calculated from Fourier transforms. To model the case of vertical orientation (Supp. Fig. 1b), the domain is orthogonal to *v*, making the RF separable for color and form – i.e. the achromatic DoG was scaled by a coefficient to create the M-opsin RF, and a different coefficient to create the S-opsin RF.

### Data analysis

*ISI imaging:* The preferred phase at each pixel, at the frequency of the drifting bar, was calculated for each of the four cardinal drift directions. The opposing directions (e.g. up and down) were then combined to yield retinotopic position as described in Kalatsky and Styker^21^. This gave a boundary of V1 that could be manually outlined.

#### Generating neuronal tuning curves from two-photon movies

Neurons were identified using the local cross-correlation image, whereby the time course of each pixel was cross-correlated with the weighted sum of its neighbors^18,49^. This is given as

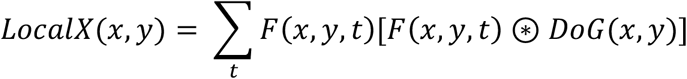

where *F*(*x, y, t*) are the fluorescence values at each pixel and timepoint, *DoG*(*x, y*) is a difference-of-Gaussian image, and ⊛ indicates convolution in *x* and *y*. The DoG rewards pixels that are correlated in time with their immediate neighborhood, but also penalizes them if they are correlated in time with the broader surround. The central Gaussian of the DoG was near the size of a neuron body (σ = 3 µm), whereas the outer “suppressive” Gaussian had σ = 20 µm to capture the surrounding neuropil. The integral of each Gaussian was normalized. Groups of pixels about the size of a cell body could be more clearly disambiguated from the surround, which is important for the subsequent manual selection of cells. The puncta were manually selected using a “point-and-click” GUI. This manual selection process was qualitatively conservative in that only the brightest puncta (i.e. most distinguishable from the surround) were selected. The min and max number of selected cells across the 16 ROIs was 46 and 165. The number selected from the ROI in Figure 2b,c was 126. After clicking on a location, a local threshold was applied to *LocalX* in order to identify the cell ROI. The pixels inside the ROI were then averaged to give the time course of each cell.

On each trial of the drifting grating stimulus, a neuron’s raw fluorescence timecourse was converted into units of Δ*F/F*. Specifically, Δ*F/F* = [*F*(*t*)-*p*]/*p*, where *F*(*t*) is the raw signal and *p* is the mean fluorescence during a blank screen preceding the trial. Gratings were presented for 3 sec. *p* was the average response over the preceding 1000 ms, and the mean of Δ*F/F* was taken between 500 ms to 3250 ms after stimulus onset. For the retinotopy stimulus, Δ*F/F* was not calculated since we were only interested in the phase.

#### Quantifying spatial frequency tuning

To get the SF tuning curve of each cell, we averaged responses over repeats, and then drift direction. Next, a Gaussian function with a domain of log_2_(SF) was fit to quantify tuning curves. The fitted parameters were μ, σ, and amplitude. The baseline was set to zero.

The peak SF and bandpass factor (Figs. 4 and 5) were taken from the fit. The peak location of the Gaussian fit differs from the parameter μ when the tuning is low-pass, or if the tuning appears high-pass by our limited SF domain. Bandpass factor was a function of the response magnitude at the peak SF (*R*_*peak*_), along with the lowest SF of 1/80 cyc/deg (*R*_*1/80*_). Bandpass factor equals (*R*_*peak*_ *-R*_*1/80*_)/*(R*_*peak*_ *+ R*_*1/80*_). If the peak is at 1/80 cyc/deg, the bandpass factor equals zero. If *R*_*1/80*_ *=* 0, it equals 1.

#### Quantifying the dependence of preferred orientation on opponency

The sampling of orientation in our stimulus was spaced by 45°. Supplemental Figure 1c compared color-form inseparability against the deviation of preferred orientation (*θ*_*pref*_) from horizontal. *θ*_*pref*_ = [∠ ∑_*θ*_ *T*(*θ*) *e*^*i*2*θ*^]/ 2, where *T*(*θ*) is the orientation tuning curve. S+R contrast was used for *T*(*θ*), and responses were averaged over repeats and SF. Opponency was quantified using a metric for the inseparability of color and form - *log*(peak_S+R_/peak_S-R_) - which can also be visualized as distance from the unity line in Figure 4i.

#### Two-photon data yield

13 animals and 16 regions-of-interest were used in the data analysis for this study. There was an additional cohort of approximately six mice in which imaging was attempted but clear neuronal responses to drifting gratings were not observed. The failure to illicit responses in some mice may be attributed to poor quality of injections and/or window implant.

From the 16 included regions-of-interest, we first extracted time courses and tuning curves using the procedures described in the sections above. Next, each neuron went through a pipeline for quality control. To be included in the analysis, each neuron had to exhibit tuning for retinotopy and SF. If the Gaussian fit to SF tuning accounted for over 70% of the variance, it was used to estimate SF tuning parameters. We required that the SF tuning along all the four axes of opsin contrast pass this criterion, which was 30% of the population. For retinotopy, we used a metric for the normalized amount of Fourier energy at the stimulus frequency. To do this, we first subtracted a linear fit to each cell’s time course to remove some of the low frequency energy. Next, we calculated the Fourier magnitude at the stimulus frequency, and divided by the sum of all Fourier magnitudes. If this fraction was below 0.2, the cell was excluded. 82% of the population passed this retinotopy criterion. Exclusion from SF tuning “OR” retinotopy gave a net yield of 385 neurons, which was 28% of the population originally obtained in the manual “point-and-click” procedure described above. These 385 neurons are represented in Fig. 2f.

Next, we excluded neurons based on their RF position and their response magnitude to the S- and R-opsin isolating color directions (Fig. 2f, dashed box). The goal of this final culling stage was to ensure that each V1 neuron was receiving input from S- and R-opsin, but little from M-opsin; interpretations of R-S opponency are less clear without this step. To calculate the balance of rod and S-opsin input, we took the peak of the SF tuning curve from R-contrast, and divided by the sum of the SF peaks from R and S contrast. This metric was defined as “%R” in Figure 2f. Of the 385 neurons that passed the quality control for SF tuning and retinotopy described above, 97% had a RF above the midline, and 37% had 1/3 < %R < 2/3. 36% passed both criteria, giving a total yield of 140 neurons. These 140 neurons are represented in Figures 3, 4, and 5. We note that the data set before (n=385) and after (n=140) culling by the dashed box in Figure 2f are provided on-line, and both give similar results. See the Discussion for additional information on how including the imbalanced population (i.e. %R near 0 or 1) affects key results.

## ACKNOWLEDGEMENTS

This work was supported by the Whitehall Foundation and the NIH R01EY028657. We are grateful to Gabriela Coello-Reyes for performing surgeries and intrinsic imaging.

## DATA AVAILABILITY

Data files for Figures 2-5 are provided as a MATLAB (.mat) file on Dryad. Included is the data before and after the culling described in Figure 2f and the Methods. Raw two-photon images are not provided due to size limitations. Also included is a ReadMe file.

## References

1. Szatko, K. P. et al. Neural circuits in the mouse retina support color vision in the upper visual field. Nature Communications 11, 3481 (2020).

2. Joesch, M. & Meister, M. A neuronal circuit for colour vision based on rod-cone opponency. Nature 532, 236–239 (2016).

3. Nadal-Nicolás, F. M. et al. True S-cones are concentrated in the ventral mouse retina and wired for color detection in the upper visual field. eLife 9,.

4. Denman, D. J. et al. Mouse color and wavelength-specific luminance contrast sensitivity are non-uniform across visual space. eLife 7, e31209 (2018).

5. Qiu, Y. et al. Natural environment statistics in the upper and lower visual field are reflected in mouse retinal specializations. Current Biology (2021) doi:10.1016/j.cub.2021.05.017.

6. De Franceschi, G., Vivattanasarn, T., Saleem, A. B. & Solomon, S. G. Vision Guides Selection of Freeze or Flight Defense Strategies in Mice. Curr Biol 26, 2150–2154 (2016).

7. Wallace, D. J. et al. Rats maintain an overhead binocular field at the expense of constant fusion. Nature 498, 65–69 (2013).

8. Yilmaz, M. & Meister, M. Rapid innate defensive responses of mice to looming visual stimuli. Curr Biol 23, (2013).

9. Hubel, D. H. & Wiesel, T. N. Receptive fields of single neurones in the cat’s striate cortex. J Physiol 148, 574–591 (1959).

10. De Valois, R. L., Albrecht, D. G. & Thorell, L. G. Spatial frequency selectivity of cells in macaque visual cortex. Vision Res. 22, 545–559 (1982).

11. Niell, C. M. & Stryker, M. P. Highly selective receptive fields in mouse visual cortex. J. Neurosci. 28, 7520–7536 (2008).

12. Solomon, S. G. & Lennie, P. The machinery of colour vision. Nat. Rev. Neurosci. 8, 276–286 (2007).

13. Johnson, E. N., Hawken, M. J. & Shapley, R. The spatial transformation of color in the primary visual cortex of the macaque monkey. Nat Neurosci 4, 409–416 (2001).

14. Ts’o, D. Y. & Gilbert, C. D. The organization of chromatic and spatial interactions in the primate striate cortex. J. Neurosci. 8, 1712–1727 (1988).

15. Lennie, P., Krauskopf, J. & Sclar, G. Chromatic mechanisms in striate cortex of macaque. J Neurosci 10, 649–669 (1990).

16. Livingstone, M. S. & Hubel, D. H. Anatomy and physiology of a color system in the primate visual cortex. J. Neurosci. 4, 309–356 (1984).

17. Wool, L. E. et al. Nonselective Wiring Accounts for Red-Green Opponency in Midget Ganglion Cells of the Primate Retina. J. Neurosci. 38, 1520–1540 (2018).

18. Rhim, I., Coello-Reyes, G. & Nauhaus, I. Variations in photoreceptor throughput to mouse visual cortex and the unique effects on tuning. Sci Rep 11, 11937 (2021).

19. Chen, T.-W. et al. Ultrasensitive fluorescent proteins for imaging neuronal activity. Nature 499, 295–300 (2013).

20. Marshel, J. H., Garrett, M. E., Nauhaus, I. & Callaway, E. M. Functional specialization of seven mouse visual cortical areas. Neuron 72, 1040–1054 (2011).

21. Kalatsky, V. A. & Stryker, M. P. New paradigm for optical imaging: temporally encoded maps of intrinsic signal. Neuron 38, 529–545 (2003).

22. Schluppeck, D. & Engel, S. A. Color opponent neurons in V1: a review and model reconciling results from imaging and single-unit recording. J Vis 2, 480–492 (2002).

23. Livingstone, M. & Hubel, D. Segregation of form, color, movement, and depth: anatomy, physiology, and perception. Science 240, 740–749 (1988).

24. Hartigan, J. A. & Hartigan, P. M. The Dip Test of Unimodality. The Annals of Statistics 13, 70–84 (1985).

25. Mechler, F. & Ringach, D. L. On the classification of simple and complex cells. Vision Research 42, 1017–1033 (2002).

26. Field, G. D. et al. Functional connectivity in the retina at the resolution of photoreceptors. Nature 467, 673–677 (2010).

27. Spitschan, M., Aguirre, G. K., Brainard, D. H. & Sweeney, A. M. Variation of outdoor illumination as a function of solar elevation and light pollution. Sci Rep 6, 26756 (2016).

28. Gouras, P. & Ekesten, B. Why do mice have ultra-violet vision? Experimental Eye Research 79, 887–892 (2004).

29. Geng, Y. et al. Optical properties of the mouse eye. Biomed. Opt. Express, BOE 2, 717–738 (2011).

30. Strasburger, H., Bach, M. & Heinrich, S. P. Blur Unblurred—A Mini Tutorial. Iperception 9, 2041669518765850 (2018).

31. Li, H., Liu, W. & Zhang, H. F. Investigating the influence of chromatic aberration and optical illumination bandwidth on fundus imaging in rats. JBO 20, 106010 (2015).

32. Girman, S. V., Sauvé, Y. & Lund, R. D. Receptive Field Properties of Single Neurons in Rat Primary Visual Cortex. Journal of Neurophysiology 82, 301–311 (1999).

33. Szél, A. et al. Unique topographic separation of two spectral classes of cones in the mouse retina. J. Comp. Neurol. 325, 327–342 (1992).

34. Chang, L., Breuninger, T. & Euler, T. Chromatic coding from cone-type unselective circuits in the mouse retina. Neuron 77, 559–571 (2013).

35. Adelson, E. H. & Bergen, J. R. Spatiotemporal energy models for the perception of motion. J Opt Soc Am A 2, 284–299 (1985).

36. Shapley, R. & Hawken, M. Color in the Cortex—single- and double-opponent cells. Vision Res 51, 701–717 (2011).

37. Wang, Q. & Burkhalter, A. Area map of mouse visual cortex. J. Comp. Neurol. 502, 339–357 (2007).

38. Garrett, M. E., Nauhaus, I., Marshel, J. H. & Callaway, E. M. Topography and areal organization of mouse visual cortex. J. Neurosci. 34, 12587–12600 (2014).

39. Martinez, L. M. et al. Receptive field structure varies with layer in the primary visual cortex. Nat. Neurosci. 8, 372–379 (2005).

40. Juavinett, A. L., Nauhaus, I., Garrett, M. E., Zhuang, J. & Callaway, E. M. Automated identification of mouse visual areas with intrinsic signal imaging. Nat Protoc 12, 32–43 (2017).

41. Rhim, I., Coello-Reyes, G., Ko, H.-K. & Nauhaus, I. Maps of cone opsin input to mouse V1 and higher visual areas. J. Neurophysiol. jn.00849.2016 (2017) doi:10.1152/jn.00849.2016.

42. Chen, T.-W. et al. Ultrasensitive fluorescent proteins for imaging neuronal activity. Nature 499, 295–300 (2013).

43. Brainard, D. H. The Psychophysics Toolbox. Spat Vis 10, 433–436 (1997).

44. Pelli, D. G. The VideoToolbox software for visual psychophysics: transforming numbers into movies. Spat Vis 10, 437–442 (1997).

45. Lei, B. & Yao, G. Spectral attenuation of the mouse, rat, pig and human lenses from wavelengths 360nm to 1020nm. Experimental Eye Research 83, 610–614 (2006).

46. Remtulla, S. & Hallett, P. E. A schematic eye for the mouse, and comparisons with the rat. Vision Res. 25, 21–31 (1985).

47. Govardovskii, V. I., Fyhrquist, N., Reuter, T., Kuzmin, D. G. & Donner, K. In search of the visual pigment template. Vis. Neurosci. 17, 509–528 (2000).

48. Naarendorp, F. et al. Dark light, rod saturation, and the absolute and incremental sensitivity of mouse cone vision. J. Neurosci. 30, 12495–12507 (2010).

49. Smith, S. L. & Häusser, M. Parallel processing of visual space by neighboring neurons in mouse visual cortex. Nat Neurosci 13, 1144–1149 (2010).

