## Supplemental material for "Joint representations of color and form in mouse visual cortex described by random pooling from rods and cones"

**Color has a weak dependence on orientation preference due to the cone-opsin gradient**

To test for a link between opsin opponency in V1, and the M-to-S opsin gradient in the retina, we performed we performed two tests. The first compared opsin opponency to RF position, with the prediction that opponency is greater near the horizontal midline since where the gradient is presumably strongest. The second compared opsin opponency to orientation preference, with the prediction that opsin opponency is stronger for neurons tuned to horizontal, since the ON and OFF channels will be unique in this case. For both tests, we used a single metric for the magnitude of opsin opponency to simplify the analyses - the shift in SF tuning between S-R and S+R contrast, which we will refer to as "inseparability". For each data point in Fig. 4i ($peak_{S-R}$ vs. $peak_{S+R}$) of the main text, inseparability is proportional to its distance from the unity line. To be clear, single-opponent neurons have the strongest opponency and the strongest inseparability in this domain.

The prediction of the first test is that that M-S opponency, and thus inseparability, should be greater at lower retinotopic position which is near the midline. All the neurons had RFs above the midline, so retinotopy in this case is effectively distance from the midline. There was not a significant correlation between retinotopy and inseparability ($p = 0.5$).

Next, we compared inseparability against preferred orientation (Supplemental Data Fig. 1c). Interestingly, this yielded the expected trend from the cone-opsin gradient ($r = -0.25$; $p = 0.002$); neurons tuned closer to horizontal were more inseparable. However, the steepness of the trend is quite shallow, and shows that the inseparability of color and form shown in Fig. 4i cannot be accounted for by the cone opsin gradient. This can most easily be explained by considering neurons tuned near vertical (right side of the scatter plot), which were still inseparable (mean above zero), and only slightly less so than the total population. Specifically, we took the subpopulation with a preferred orientation less than 30° from vertical ($n = 36$), and calculated the same statistics shown in the inset of Fig. 4i, which gave $\Delta = 41\%$, $p = 1.8 \times 10^{-6}$. To summarize, the analyses reveal that there is indeed likely to be some opponency caused by the cone-opsin gradient in the mesopic regime, but it is minimal compared to the rod-cone opponency.

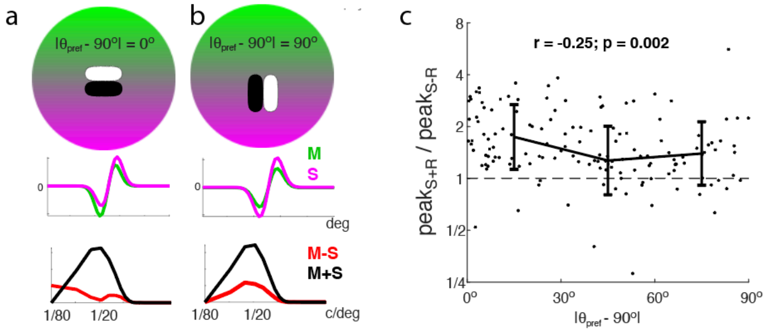


**Supplemental Data Figure 1: A dependence of inseparability on orientation preference predicts marginal opponency from the cone-opsin gradient in the mesopic state.** (a) The colored disk is a cartoon representation of the cone-opsin mosaic, where green and violet are M and S-opsin expression. For the actual simulation, we used a model of this gradient based on retinal ganglion cell output (ref. Wang et al. 2011). Overlaid at the steepest part of the gradient is a cartoon V1 RF tuned for horizontal orientation ($\theta_{pref} = 90^o$). The RF is composed of an ON and OFF Gaussian that generates a preferred SF of 0.05 cyc/deg. The green and violet traces below are the 1D profiles of a simulated S and M-opsin RF, calculated by multiplying the monochromatic spatial RF by the percentage of S and M-opsin expression at each location given by the model in Wang et al 2011. The red and black traces on bottom are the corresponding SF tuning curves for M-S and M+S contrast. This RF has lowpass tuning for M-S contrast and bandpass tuning for M+S contrast, making it inseparable for color and form. (b) Same as 'a' but the RF prefers vertical orientation and is positioned more ventral. This RF has the same bandpass tuning for M-S and M+S contrast, making it separable for color and form. (c) Actual data showing color-form inseparability vs. orientation preference. The y-axis values can be calculated from the scatter plot values in Figure 4i. The x-axis is preferred orientation, relative to horizontal ($90^o$). Error bars show the median and SD in three bins, $0^o$-to-$30^o$, $30^o$-to-$60^o$, and $60^o$-to-$90^o$.
